# Comparative Analysis of Rumen Microbial Communities in Japanese Black Cattle: Site and Fertility Influences on Microbiome Composition and Function

**DOI:** 10.1101/2025.10.16.682985

**Authors:** Vincent Pamugas Reyes, Yukihiro Umeda, Sho Nakamura, Yasuhiro Morita, Shuichi Matsuyama, Shunsaku Nishiuchi, Jun Murase, Satoshi Ohkura

## Abstract

**Introduction:** The rumen microbiome plays a crucial role in nutrition, and productivity of cattle, influencing processes such as digestion, metabolism, and overall health. Understanding the composition and diversity of microbial communities within the rumen is essential for improving cattle management and enhancing the overall health outcomes. This study aims to investigate the rumen microbiome composition across two farms and fertility levels in Japanese Black beef cattle, using 16S rRNA amplicon sequencing. Differences in functional pathways between low- and normal-fertility cattle were also investigated.

**Result:** Core ruminal microbes were identified with *Bacteroidota* being predominant across sites and fertility levels. Alpha and beta diversity metrics showed that the site explained a substantial variation in microbiome composition, while fertility had a minimal impact. Differential abundance analysis using LEfSe identified distinct microbial profiles for each site. Notably, taxa such as *Fibrobacterota* and *Negativicutes* were more abundant on one site, whereas *Acholeplasmataceae* and *Lentisphaeria* were more prevalent on the other site

**Conclusion:** Using the sparse Partial Least Squares Discriminant Analysis (sPLS-DA) key taxa associated with fertility status, including *Bradymonadales* and *Elusimicrobiaceae* for low fertility and *Anaerovoracaceae* and *Desulfovibrionaceae* for normal fertility were identified. In addition, enhanced pathways in the normal fertility group include nicotinate degradation and complex sugar catabolism, while the low-fertility group showed increased activity in glycine betaine and nitrate reduction pathways, suggesting metabolic shifts and may impact reproductive efficiency. These findings underscore the complex interplay between geographic and biological factors in shaping the rumen microbiome.

## Introduction

The complex ecosystem of the rumen, a vital compartment of the ruminant digestive system, harbors a dense and diverse microbiota. This microbial community, predominantly composed of bacteria, plays a pivotal role in fermenting complex plant materials, producing volatile fatty acids essential for energy, synthesizing vitamins and amino acids, and protecting against pathogens. In domesticated farm animals such as cattle, the ruminal microbiome’s composition and functionality directly influence feed efficiency and overall health (Li et al., 2019; Takizawa et al., 2023).

Understanding the dynamics and diversity of the ruminal bacterial community in cattle is crucial for ensuring sustainable production practices. Among these cattle, the Japanese Black breed holds significant economic importance for meat production in Japan. Focused studies on this breed are particularly relevant as reproduction emerges as a critical trait for the industry’s success (Hirooka, 2014). The studies on artificial insemination (AI) in Japanese Black beef cattle have provided detailed insights into the factors that influence reproductive efficiency. For example, the study conducted by Sasaki et al. (2016) have illustrated how season, parity, and herd size affect reproductive performance in intensively reared Japanese Black beef cattle, finding that calving and AI during winter and spring, higher parity, and smaller herd size were linked to decreased reproductive performance. Additionally, challenges like cold environmental conditions and the frequency of AI services have been explored to determine their effects on conception rates and overall reproductive health (Irikura et al., 2018; Nabenishi & Yamazaki, 2017). Recent research has highlighted the importance of microbial communities in the gastrointestinal tract and their potential impact on various health parameters, including fertility (Ashonibare et al., 2024; Kobayashi, 2023; Lehtoranta et al., 2022; Li et al., 2019). The microbiota of the female reproductive tract, particularly the vagina, has been well-documented for its role in reproductive health. However, the concept of an extra-vaginal microbiota and its influence on fertility is an emerging area of interest (Garcia-Garcia et al., 2022). Recent studies have suggested that the gut and oral microbiota may also play significant roles in reproductive events (Comizzoli et al., 2021; Ji et al., 2019; O’Callaghan et al., 2016; Xu et al., 2020). The rumen, as a major microbial habitat in ruminants, could similarly influence reproductive health making it an important area for exploring potential links to reproductive health.

In this study, we aim to: (i) characterize the composition and structure of the ruminal bacterial community in Japanese Black beef Cattle, highlighting the presence of core taxa across the select farms in Aichi Prefecture and fertility levels; (ii) investigate the associations between the ruminal microbiome profiles, geographical location of the farm, and fertility levels; (iii) identify the functional prediction of the rumen microbial communities in low and normal fertility cattle.

## 1 Materials and Methods

### 1.1 Animal and Farm Management

The present study was conducted at either a farm (Togo Farm) of the Field Science Center, Graduate School of Bioagricultural Sciences, Nagoya University in Togo-town, or a commercial farm (Oguri Farm) located in Handa City, Aichi, Japan. Both farms are 28.7 km apart in a straight line. The animals were kept in groups in a free barn and were fed mixed feeds at 0900 h and 1600 h with free access to water and licking minerals. The diet was formulated to meet or exceed nutritional requirements for non-lactating Japanese Black cows (Japanese Feeding Standard for Beef Cattle, 2008). All experimental procedures were approved by the Committee of the Care and Use of Experimental Animals at the Graduate School of Bioagricultural Sciences, Nagoya University.

A total of 32 cattle were included in the study, 19 cattle were kept at the Togo Farm and the remaining 13 cattle were kept at the Oguri Farm, categorized based on their fertility status and exposure to AI. Six cattle were classified as having low fertility, having been subjected to AI three times without conceiving. Nineteen cattle were categorized as having normal fertility, having successfully conceived after three AI attempts. The remaining seven cattle had not been subjected to AI at all.

### 1.2 Sample Collection, DNA extraction, and Sequencing

The rumen fluid collection and processing protocol were based from Sato et al. (2021). Briefly, rumen fluid samples were obtained via stomach tubing method from each animal. The initial sample was discarded to prevent saliva contamination, followed by the collection of a new sample. Then after, the rumen samples were filtered using four-layered cheesecloth. The filtered rumen samples were then stored in a −80°C freezer.

DNA was extracted from the thawed rumen samples using the SPINeasy DNA Pro Kit for Soil (MP Biomedicals, Santa Ana, CA, USA) and the FastPrep-24TM classic bead beating grinder and lysis system (MP Biomedicals). Briefly, 500 µL of each sample was processed for DNA extraction according to the manufacturer’s instructions, which included cell lysis and DNA purification. The integrity and concentration of the DNA were evaluated using gel electrophoresis and a NanoDrop spectrophotometer, respectively. Subsequently, the DNA samples were forwarded to the Beijing Genomics Institute (BGI) for sequencing, specifically targeting the V3-V4 variable regions of the 16S rRNA gene.

### 1.3 Bioinformatics Analytical Pipeline

The obtained pre-cleaned reads from the sequencing company were further processed using the <u>Q</u>uantitative Insights Into <u>M</u>icrobial <u>E</u>cology 2 (QIIME2) (Bolyen et al., 2019). The <u>D</u>ivisive <u>A</u>mplicon <u>D</u>enoising Algorithm 2 (DADA2) (Callahan et al., 2016) function from QIIME2 was used to further clean the data. In brief, reads were truncated on a quality threshold of 5 per read. The algorithm learned from 1,000,000 reads to establish reliable error models, and the ‘consensus’ method was used to detect and remove chimeric sequence. Quality-controlled ASVs were classified using the classify consensus-vsearch method against the recent version of SILVA database (version 138-99) (Quast et al., 2012) with settings including: (i) 90% identity threshold, (ii) 0.8 query coverage, (iii) and a maximum of 7 accepted hits. Further data analyses and statistical comparisons were made using R and the following R-packages: “phyloseq”, “vegan”, “MicrobiotaProcess”, “coin”, and “mixOmics” (Hothorn et al., 2008; McMurdie & Holmes, 2013; Rohart et al., 2017; S. Xu et al., 2023). The analyses also employed the following general R-packages: “dplyr”, “ggplot2”, “tidyverse”, and “VennDiagram”, and “tayloRswift” (Chen, 2022; Stephenson, 2021; Wickham et al., 2019; William et al., 2023)

### 1.4 Statistical analyses

To assess the alpha diversity of the rumen microbiome, samples were first subjected to rarefaction to equalize sequencing depth across all samples. This was achieved using the rarefy_even_depth function from the phyloseq package in R. The rarefaction depth was set to 90% of the minimum sample size, ensuring a consistent comparison base without replacement. Following rarefaction, alpha diversity indices, including the Shannon diversity index and observed features (*ASVs)*, were calculated using the estimate_richness function from the same package. Differences were statistically analyzed using the Wilcoxon test. Beta diversity was evaluated utilizing Bray-Curtis and unweighted UniFrac dissimilarity metrics. In addition, permutational multivariate analysis of variance (PERMANOVA) using the ‘adonis2’ function of the ‘vegan’ R package, with 999 permutations, was performed.

In this study, we conducted a differential abundance analysis to explore variations in microbial communities between the two sites, utilizing a modified LEfSe approach included in a custom script applied to a phyloseq object. The initial comparative analysis involved the Kruskal-Wallis test to determine if the distributions of microbial taxa differed significantly across groups defined by ’Site’. Only taxa with a p-value less than 0.01 were advanced for further analysis under strict filtering criteria. The subsequent pairwise comparisons were performed using the Wilcoxon rank-sum test. The effect size estimation was conducted through Linear Discriminant Analysis (LDA) with a score threshold of 3, highlighting features with notable differences in abundance. Both stages of comparative analysis maintained a stringent significance level of 0.01 (firstalpha and secondalpha), ensuring that the detected differences were statistically robust and biologically meaningful.

The <u>s</u>parse <u>P</u>artial <u>L</u>east <u>S</u>quares <u>D</u>iscriminant <u>A</u>nalysis (sPLS-DA) was used to identify key microbial taxa associated with fertility levels. The analysis began with principal component analysis (PCA) to explore variance of ASVs. To refine the analysis, an initial sPLS-DA model was constructed with 10 components, visualized through projection plots. Performance evaluation of the initial sPLS-DA model, using 5-fold cross-validation repeated 50 times, determined the optimal number of components. This was achieved by assessing various model configurations for balanced error rate (BER) and AUC values. The performance assessment indicated the optimal model configuration. Subsequent tuning identified the ideal number of variables for each component, leading to the final optimized sPLS-DA model. Associations between microbial taxa and fertility status were further explored using the final sPLS-DA model.

Functional pathways were analyzed using PICRUSt2-processed 16S rRNA gene amplicon sequencing data. Pathway abundances were normalized for sequencing depth variations. The methodology for developing initial and final sPLS-DA models for pathway analysis mirrored the approach taken for microbial taxa. The final sPLS-DA model loadings were determined using the ’mean’ and ’max’ contrib methods to highlight the most influential pathways.

## 2 Result

### 2.1 Amplicon sequence variants (ASVs) and taxa profiling

In the analysis of the rumen microbiome, site and fertility level were considered as factors. All 32 samples were included for site comparison, whereas only samples from cattle with low and normal fertility were considered for fertility analysis, excluding the unexposed group due to their lack of relevance to fertility outcomes.

The comparative analysis of ASVs using Venn diagrams illustrated significant overlaps and unique ASVs between different sites and fertility levels (Figures 1A and 1B). For site comparisons, the Oguri and Togo samples showed a substantial overlap, with 34% (4,962) of the ASVs shared between them. Each site also maintained a significant portion of unique ASVs, with Oguri having 33% (4,739) and Togo having 33% (4,863) unique ASVs. In the analysis of fertility levels, intersections among the normal and low fertility groups revealed both distinct and shared ASVs. The normal group had 5,325 (41%) unique ASVs, while the low group had 1,135 (9%). Notably, a significant overlap of 50% (6,556) of the ASVs was observed between the normal and low fertility groups.

**Figure 1.**
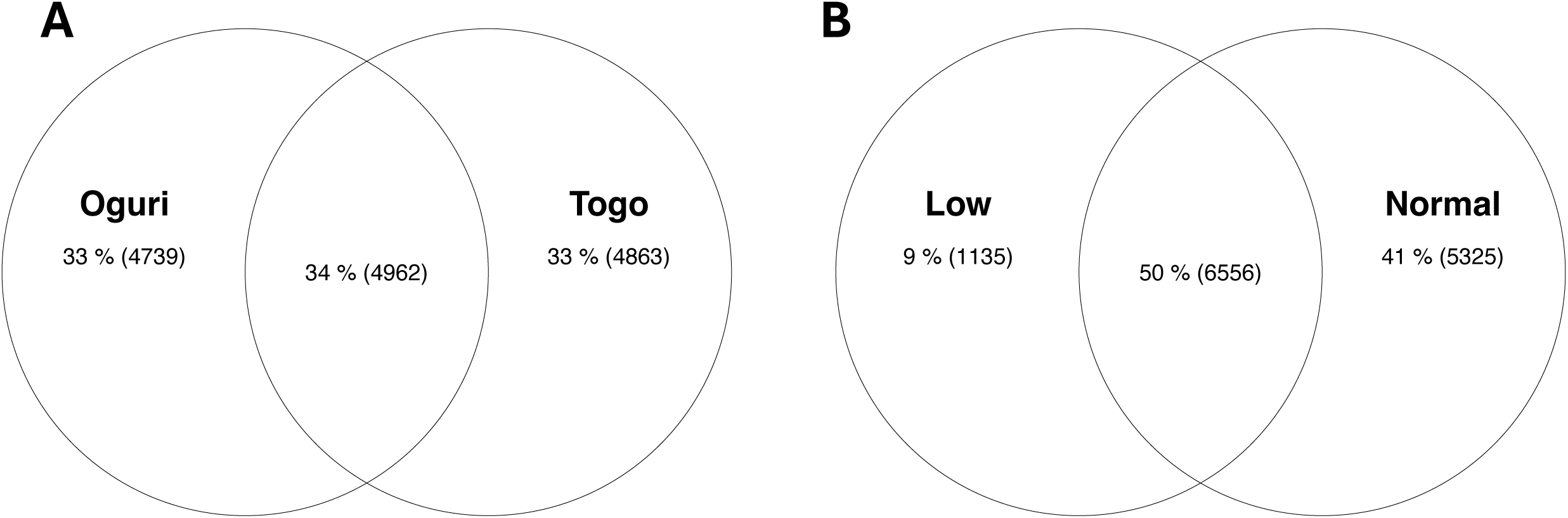
Comparative analysis of amplicon sequence variants (ASVs) across: (A) Sites-Venn diagram depicting the shared and unique ASVs between Oguri and Togo sites; (B) Venn diagram showing the distribution of ASVs among fertility groups: normal and low

Core ruminal microbes were identified at the phylum and family levels by filtering the dataset to include only ASVs detected at or above the threshold in at least 95% of samples. The data of core taxa was summarized in Supplementary Tables 1-4. The analysis of the relative abundance of bacterial phyla in the rumen microbiome from cattle at Oguri and Togo, as well as among different fertility levels, revealed a consistent pattern (Figures 2A and 2B). Both comparisons—across the two sites and between fertility groups—demonstrated a predominant presence of *Bacteroidota* and *Firmicutes*. Additionally, slight variations were observed in less abundant phyla like *Fibrobacterota* and *Patescibacteria* at both sites, with minor differences in *Patescibacteria*, *Verrucomicrobiota*, and *Fibrobacterota* among the fertility groups.

**Figure 2.**
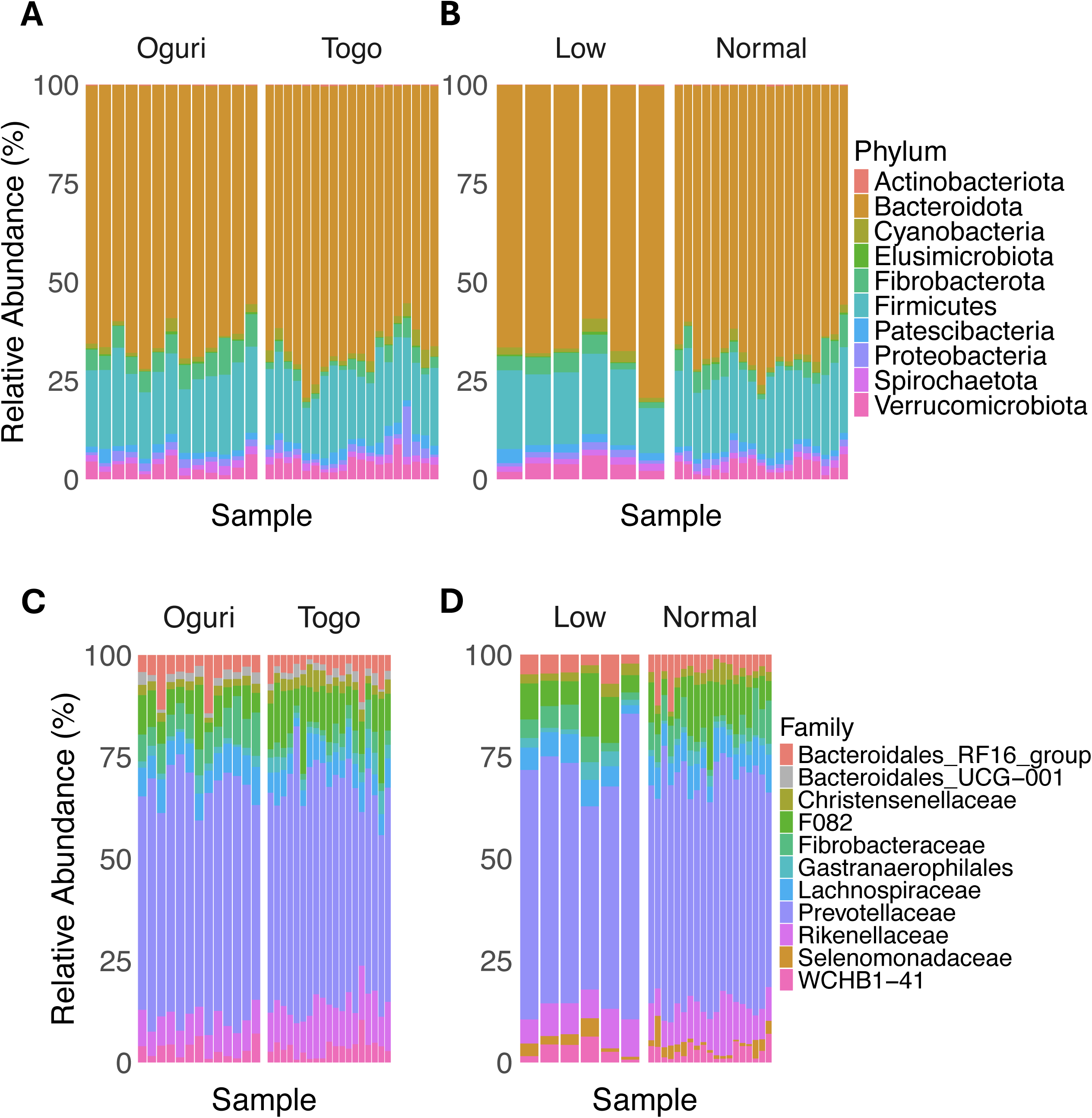
Stacked bar plots depicting the composition of microbial communities in Oguri and Togo, and in low and normal fertility groups. (A) Microbial community composition at phylum level by site. (B) Microbial community composition at phylum level by fertility status. (C) Microbial community composition at family level by site. (D) Microbial community composition at family level by fertility status.

At the family level, the analysis of the relative abundance of bacterial taxa in the rumen microbiome from cattle at Oguri and Togo, as well as among different fertility levels, showed similar patterns (Figure 2C and 2D). Across both sites, *Prevotellaceae*, *Bacteroidales*_RF16_group, and *Rikenellaceae* were the most abundant families, with slight variations in *Lachnospiraceae* and *Fibrobacteraceae*. Similarly, the comparison between low and normal fertility levels in cattle revealed a consistent dominance of *Prevotellaceae*, followed by *Rikenellaceae* and *Bacteroidales*_RF16_group, with minor variations in *Lachnospiraceae* and *Fibrobacteraceae*.

### 2.2 Diversity analysis of rumen microbiome

We explored the impact of site and fertility levels on microbial diversity and community structure. Initially, we assessed alpha diversity using the Shannon diversity index and the observed ASVs (Figures 3A and Fig 3B). The Wilcoxon test revealed a significant difference in Shannon diversity index values between the Oguri and Togo sites (Figure 3B). Specifically, Oguri exhibited a significantly higher Shannon index compared to Togo (*P < 0.05*), indicating greater microbial diversity. In contrast, the comparison of observed ASVs between the two sites showed no significant differences, confirming that the total number of distinct microbial species present at each site was comparable. Next, we examined the influence of fertility levels on microbial diversity, specifically looking at the Shannon Index and the observed ASVs (Figure 3A). The Wilcoxon test applied to these data showed no significant differences across the two fertility levels.

**Figure 3.**
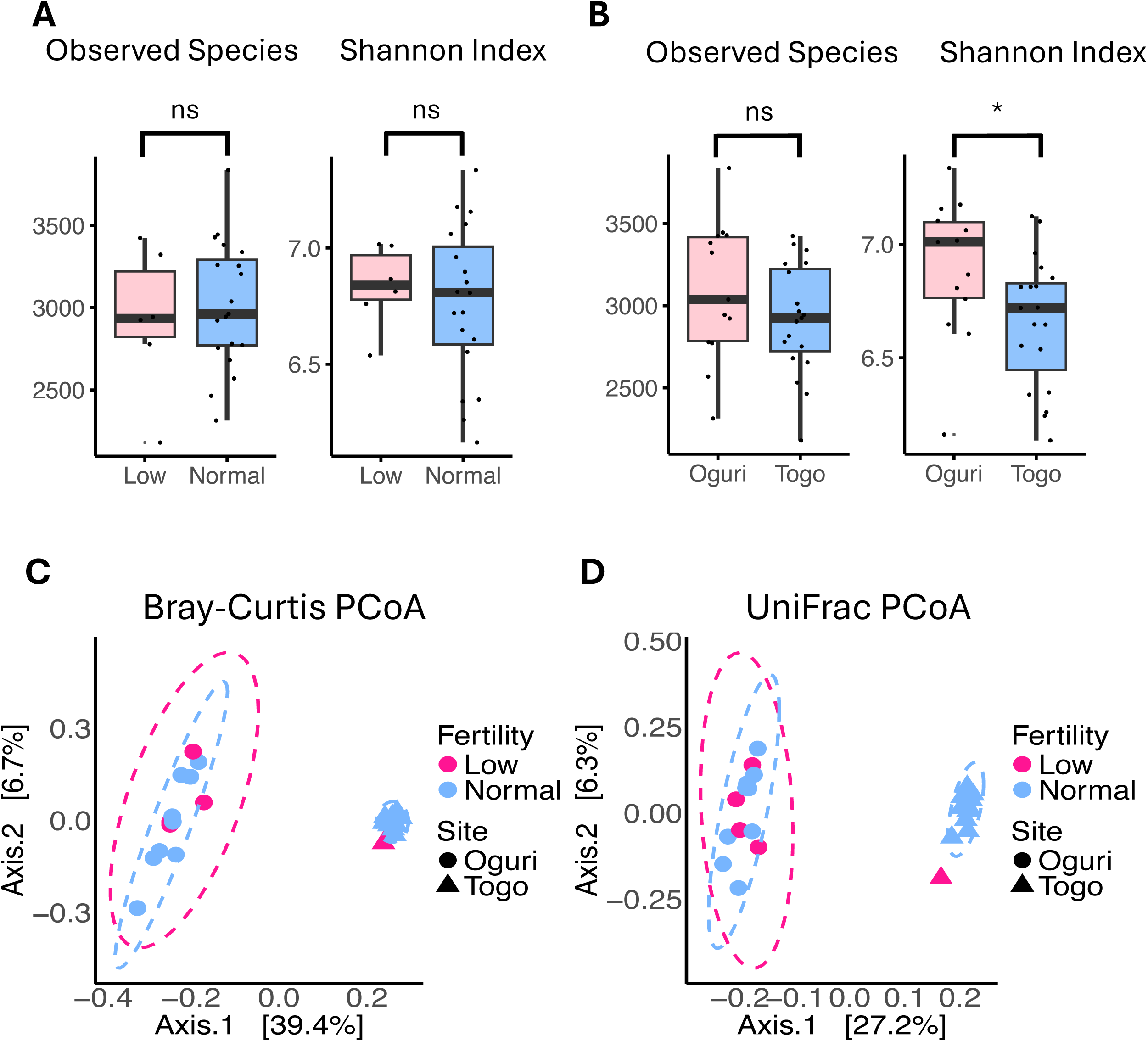
Alpha and Beta Diversity Metrics. (A) Box plots of Shannon index and observed species representing alpha diversity of the microbiome in Togo and Oguri Farms; (B) Box plots of Shannon index and observed species representing alpha diversity of the microbiome in samples with varying fertility levels; (C) Principal coordinate analysis (PCoA) based on Bray-Curtis dissimilarities on the microbial communities (ASVs level) associated with sites and fertility levels.

To further investigate the effects of fertility and site on the rumen microbiome composition, we employed beta diversity metrics, using Bray-Curtis dissimilarity and unweighted UniFrac distance. Using the Bray-Curtis dissimilarity matrix, the analysis indicated that fertility explained a small proportion of the variation in the microbiome composition (R² = 0.032, *P = 0.633*). In contrast, the effect of site explained a substantial proportion of the variation in the microbiome composition (R² = 0.389, *P = 0.001*). Similarly, using the UniFrac distance matrix, the analysis showed that fertility accounted for a small proportion of the variation in the microbiome composition (R² = 0.04028, *P = 0.381*). The effect of site, however, was significant, explaining a substantial portion of the variation in the microbiome composition (R² = 0.271, *P = 0.001*).

### 2.3 Differences in microbiota composition between sites/fertility levels

The differential abundance analysis between the microbial communities of Oguri and Togo showed variations in the relative abundances of several bacterial taxa. The relative abundance boxplots (Figure 4A) demonstrate the distribution of various bacterial taxa between Oguri and Togo. The analysis indicated that taxa such as *Rikenellaceae*, *Vampirivibrionia*, *Gastranaerophilales*, and *Cyanobacteria* were significantly more abundant in Togo. In contrast, taxa such as *Fibrobacteraceae*, *Selenomonadaceae*, *Negativicutes*, and *Acholeplasmatales*, showed higher relative abundances in Oguri. The LDA score plot (Figure 4B) further supports these findings by showing the effect sizes of the differentially abundant taxa. In Oguri, taxa including *Fibrobacterota*, *Fibrobacteria*, and *Negativicutes* displayed significantly higher LDA scores. Conversely, Togo exhibited higher LDA scores for *Rikenellaceae*, *Vampirivibrionia*, *Gastranaerophilales*, and *Cyanobacteria*.

**Figure 4.**
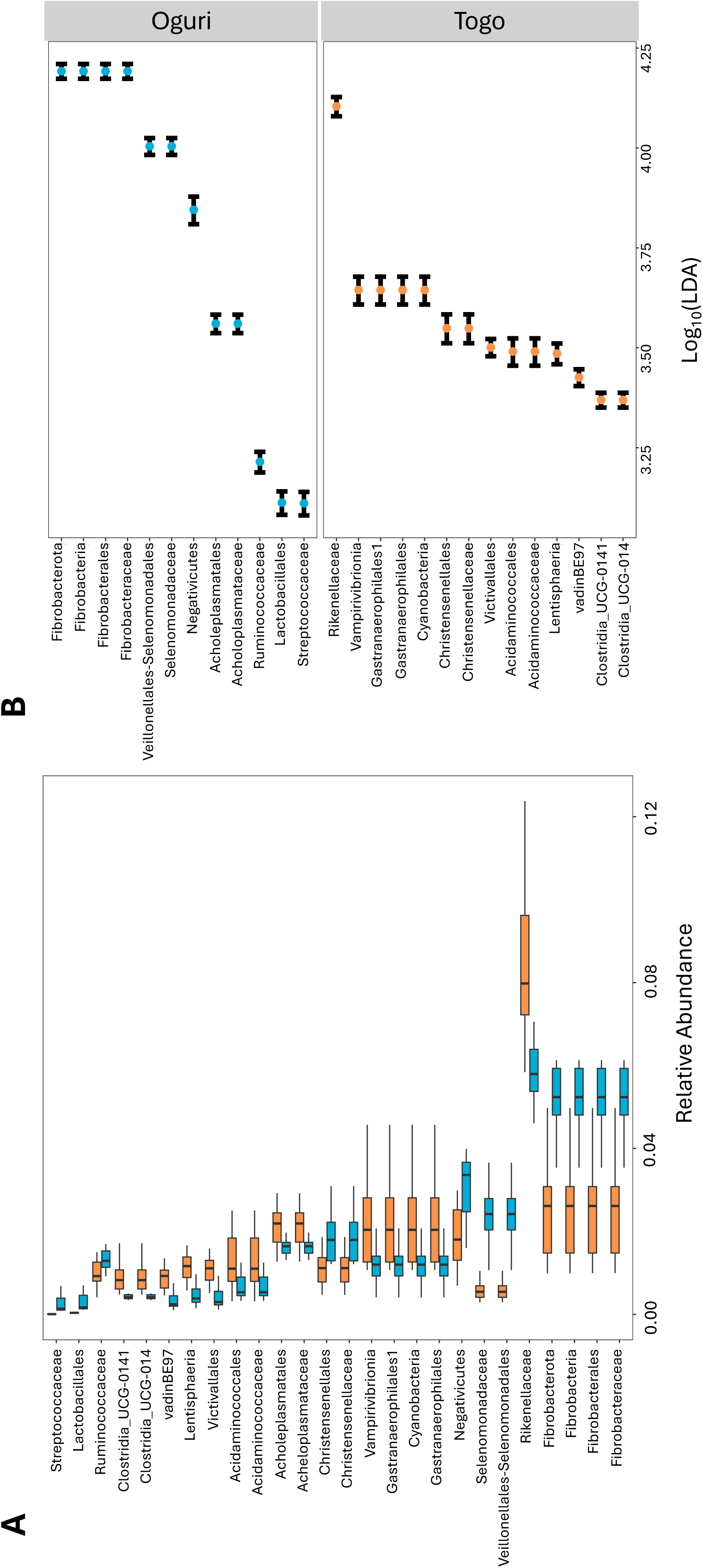
Differential abundance and effect size of microbial taxa between Oguri and Togo. (A) Boxplot showing the relative abundances of various bacterial taxa in Oguri (blue) and Togo (orange). (B) LDA score plot illustrating the effect sizes of differentially abundant taxa.

Despite the lack of distinct clustering between normal and low fertility levels in the Bray-Curtis and UniFrac PCoA (Fig 3C-3D), the application of sPLS-DA enabled the identification of key taxa unique to both low and normal fertility samples. Initially, PCA was employed as an unsupervised method to explore the variance within the data, providing a preliminary understanding of the sample distribution. The PCA revealed no distinct clustering between normal and low fertility groups (Figure 5A). The final sPLS-DA model delineated the fertility groups, confirming the presence of distinct microbial signatures associated with normal and low fertility (Figure 5B).

**Figure 5.**
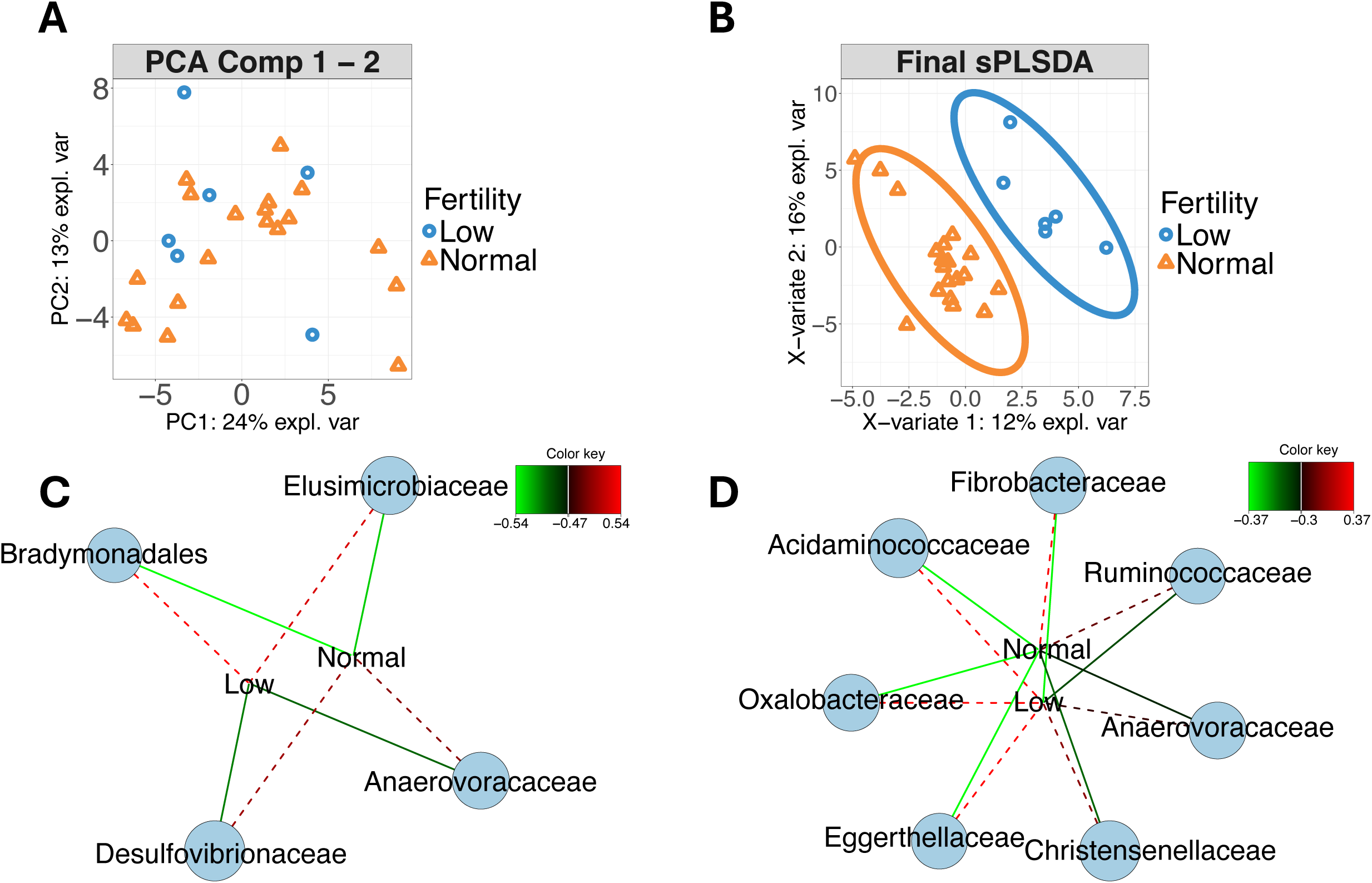
Microbial community analysis in relation to cattle fertility levels. (A) Principal Component Analysis (PCA) plot showing the samples based on fertility levels; (B) Sparse Partial Least Squares Discriminant Analysis (sPLS-DA) plot highlighting the distinct clustering of samples with normal and low fertility. (C) Network plot (Component 1) illustrating positive (dashed lines) and negative (solid lines) associations between specific microbial families and fertility levels (D) Extended network plot (Component 2) showing additional microbial families. Color key indicates the strength of associations.

Association analyses of the sPLS-DA model identified distinct microbial signatures associated with fertility status. In component 1 (Figure 5C), the taxa *Bradymonadales* and *Elusimicrobiaceae* exhibited positive associations with the low fertility group and negative associations with the normal fertility group. In contrast, *Anaerovoracaceae* and *Desulfovibrionaceae* were negatively associated with low fertility and positively associated with normal fertility. Component 2 further distinguished microbial communities (Figure 5D), revealing that *Fibriobacteraceae* and *Ruminococcaceae* had negative associations with low fertility and positive associations with normal fertility. Conversely, *Anaerovoracaceae*, *Christensenellaceae*, *Eggerthellaceae, Oxalobacteraceae*, and *Acidaminococcaceae* displayed positive associations with low fertility and negative associations with normal fertility.

### 2.4 Differences in functional pathways between low and normal fertility cattle

Functional differences between cattle with normal and low fertility were analyzed using metabolic pathways inferred from 16S rRNA gene amplicon sequencing data, processed through PICRUSt2. To adjust for variations in sequencing depth across samples, pathway abundances were normalized by converting raw counts to proportional abundances. The PCA, shown in Figure 6A, revealed that the first two principal components accounted for a substantial portion of the variance (PC1: 36% and PC2: 22%). Further refinement using sPLS-DA, as depicted in Figure 6B, emphasized the distinction between the groups, with the model explaining a total of 37% of the variance (X-variate 1: 32% and X-variate 2: 5%). This analysis pinpointed specific metabolic pathways that vary significantly between the normal and low fertility groups.

**Figure 6.**
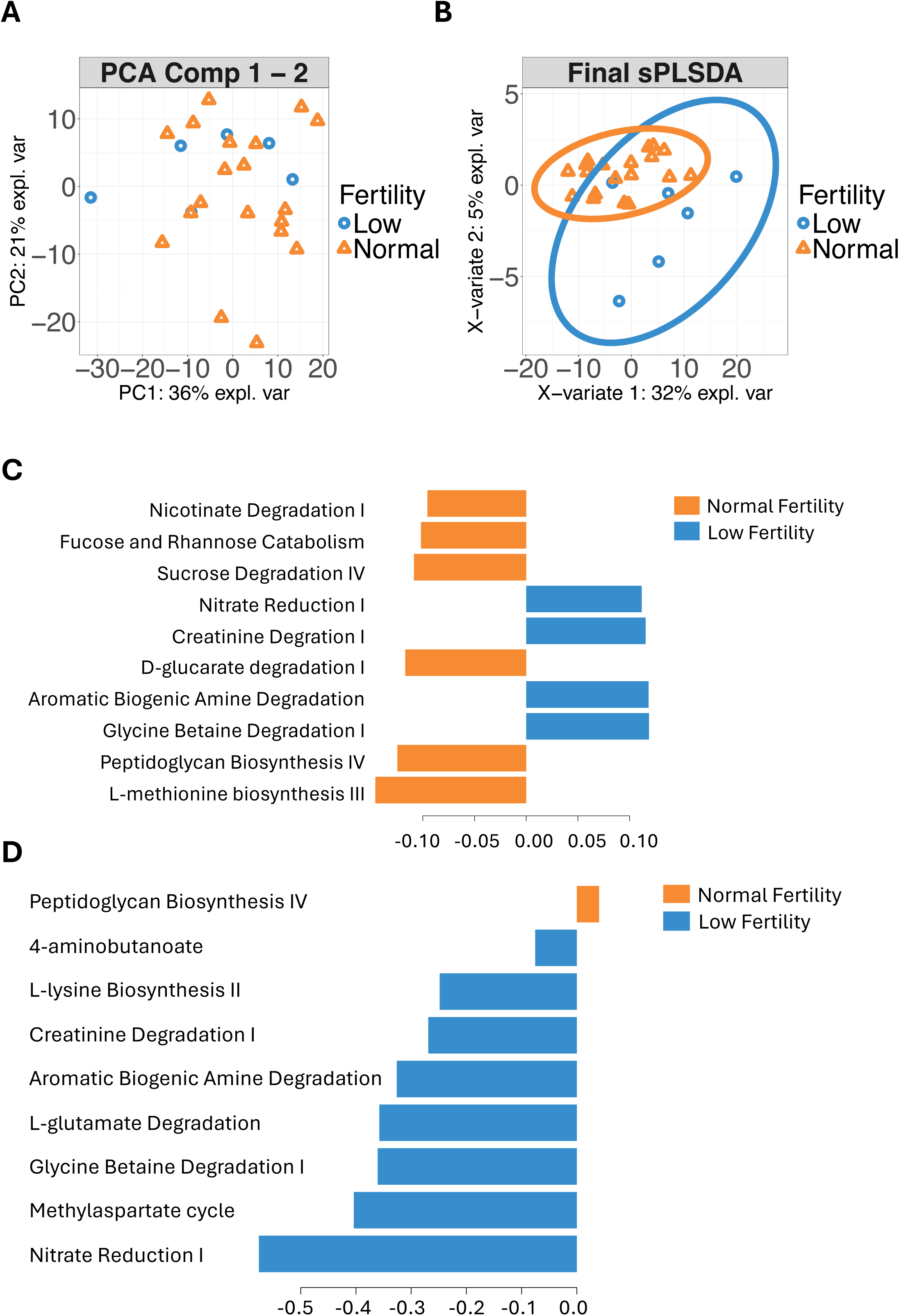
sPLS-DA Analysis Results of Pathways. (A) Principal Component Analysis (PCA) plot showing the samples based on fertility levels; (B) Final model Sparse Partial Least Squares Discriminant Analysis (sPLS-DA) plot highlighting the distinct clustering of samples with normal and low fertility. (C) Loadings Plot (First Component): Top ten metabolic pathways that have the most significant influence on the first component of the sPLS-DA model. (D) Loadings Plot (Second Component): Shows the loadings for the second component of the sPLS-DA model, highlighting the metabolic pathways that significantly influence this component.

Loadings for the sPLS-DA model were identified using the ’mean’ method to calculate the average contribution of each pathway across the model components, and the ’max’ contrib option was used to extract the pathways with the maximum impact on the model’s discrimination ability. This approach allowed for a focused analysis of the most impactful pathways, enhancing the interpretability of multivariate data. Figure 6C highlights pathways with greater activity in the normal fertility group, such as nicotinate degradation I, fucose and rhamnose catabolism, and sucrose degradation IV. These pathways are predominantly associated with vitamin B3 metabolism, complex sugar catabolism, and disaccharide metabolism, respectively.

Several pathways are more prominently represented in the low fertility group, as shown in Figure 6D. These include glycine betaine degradation I and nitrate reduction I, which are involved in the metabolism of glycine betaine and nitrate, respectively. Other pathways, such as the methylaspartate cycle and L-glutamate degradation, also show increased representation in this group. These metabolic shifts suggest that cattle with lower fertility might be experiencing changes in amino acid and nitrogen handling, which could impact their reproductive efficiency.

## 3 Discussion

The comparative analysis of rumen microbial communities in Japanese Black cattle across different fertility levels and select farms in Aichi, using 16S rRNA gene amplicon sequencing, provides crucial insights into how geographic and physiological factors shape the rumen microbiome. Previous research has emphasized the importance of the rumen microbiome in influencing host health and productivity (Fonseca et al., 2023; Li & Guan, 2017).

Our study observed substantial overlaps and unique microbial communities between sites, with a notable overlap of ASVs between Oguri and Togo, suggesting a common core of microbial taxa. The presence of unique ASVs at each site highlights the influence of specific local environmental factors on the rumen microbiome. The overlap of ASVs between normal and low fertility groups suggests a core microbial community essential for basic rumen functions.

The relative abundance of bacterial phyla in the rumen microbiome revealed notable patterns. *Bacteroidota* emerged as the predominant phylum, in the rumen microbiome followed by *Firmicutes* as the second most abundant phylum, these findings were similar with the previous studies on rumen microbiome profiling (Keum et al., 2024; Pinnell et al., 2022; Pitta et al., 2016). Despite Oguri and Togo being located in the same prefecture and at similar altitudes, subtle differences in other environmental or biotic factors might still contribute to variations in microbial abundance observed between the sites. For example, *Verrucomicrobiota* was more abundant in Togo, suggesting site-specific microbial adaptations due to differences in environmental factors and forage types available at each site (Stewart et al., 2018). Comparisons of fertility levels revealed consistent dominance of *Bacteroidota* and *Firmicutes*, regardless of fertility status, indicating these phyla’s crucial roles in rumen microbial processes. Minor variations in other phyla, such as *Verrucomicrobiota*, suggest subtle differences in microbial community composition related to fertility. At the family level, *Prevotellaceae* was the most prevalent across all samples, underscoring its ecological dominance and critical role in rumen microbial processes. Other taxonomic families showed consistent patterns across sites and fertility levels, reflecting the stable yet dynamic nature of the rumen microbiome.

While our diversity analysis revealed higher microbial diversity in the rumen microbiome of Oguri compared to Togo, these differences were not mirrored in fertility levels. This suggests that microbial diversity alone may not be a direct indicator of fertility status. This observation aligns with previous studies indicating that the functional capacities of the microbiome may be more crucial than diversity per se in influencing reproductive outcomes. For instance, Adnane & Chapwanya (2022) highlighted that the balance and specific functions of microbial communities are more directly linked to fertility outcomes than diversity alone. Similarly, Luecke et al. (2022) highlighted that the individual and interactive roles of microbial communities within the bovine microbiome are intricately linked to cattle fertility.

In dissecting the association between rumen microbiota and fertility, our results highlighted distinct microbial signatures linked to different fertility levels. Notably, taxa such as *Bradymonadales* and *Elusimicrobiaceae* were more prevalent in low fertility cattle. Similarly, a study by Yagisawa et al. (2023) analyzing the uterine microbiome identified *Bradymonadales* and its co-occurrence with *OLB8* as associated with low fertility. In addition to these taxa, we also identified other taxa that are associated metabolic pathways that were previously identified in the microbiome uterine of low fertility cattle. For example, in a study conducted by Webb et al. (2023) cattles that remained open after AI was identified to have higher relative abundance of *Christensenellaceae*. Our findings coincide with their result wherein *Christensenellaceae* in the uterus were identified to be associated to low fertility cattle groups. *Christensenellaceae* is associated with short-chain fatty acid synthesis particularly as a butyric acid producing species (Sun et al., 2024). Interestingly, butyric acid produced by bacteria has been previously identified to be involved in animal reproduction by directly regulating progesterone and 17β-Estradiol secretion (Lu et al., 2017). While such findings suggest potential as biomarkers for fertility, it is crucial to acknowledge that our study does not establish causality.

The functional analysis of metabolic pathways in low-fertility cattle revealed significant alterations, particularly in the nitrate reduction I, methylaspartate cycle, and glycine betaine degradation I pathways. These pathways are crucial to energy, carbohydrate, and amino acid metabolism, respectively. The nitrate reduction I pathway, an alternative hydrogen sink in the rumen, is linked to adapting to hypoxic conditions or changes in the microbial environment (Latham et al., 2016). Its elevated presence in low fertility cattle suggests an adaptive response to environmental stressors affecting the rumen’s microbial ecosystem. Such stressors can impact overall metabolic efficiency and reproductive systems. Environmental factors like heat, cold, humidity, and nutritional challenges influence the endocrine system and affect reproductive performance (Bova et al., 2014; Wrzecińska et al., 2021). The methylaspartate cycle, involved in assimilating acetate into central metabolism in haloarchaea, produces malate for anabolism (Borjian et al., 2016). Its higher abundance in low-fertility cattle possibly suggests a shift towards energy production from amino acid catabolism, a compensatory response to negative energy balance, which affects fertility in cattle (Sejian et al., 2012). The enhanced activity of the glycine betaine degradation I pathway aligns with its roles in metabolic adaptation to stress, where betaine acts as a stress protectant or a methyl donor (Wang et al., 2013; Zou et al., 2016). This heightened activity might suggest a compensatory response to metabolic stress or inefficiency, possibly impacting reproductive capabilities. The increased activity of this cycle may reflect an altered metabolic state where energy allocation is prioritized towards basic maintenance rather than reproductive processes. The association of these pathways with low fertility underscores the potential impact of metabolic and environmental stress on reproductive outcomes. While the protective functions of these pathways are supported by scientific literature, the direct link between these pathways’ activity and cattle fertility remains to be conclusively determined.

## 4 Conclusion

Our comprehensive analysis of the rumen microbiome in Japanese Black cattle across varied fertility levels and geographic locations underscores the significant influence of both physiological and environmental factors on microbial communities. Our study not only confirms the presence of a core microbial community essential for rumen function regardless of site and fertility differences but also reveals significant microbial signatures and metabolic pathways associated with different fertility levels. The predominance of *Bacteroidota* and *Firmicutes* across all samples, regardless of the geographic location or fertility status, highlights their fundamental role in the rumen ecosystem.

However, the identification of distinct taxa in low fertility cattle and the association of specific metabolic pathways such as Nitrate Reduction I and Glycine Betaine Degradation I with fertility levels suggest potential targets for improving reproductive outcomes. These results provide a foundation for future research aimed at elucidating the direct mechanisms by which the rumen microbiome influences fertility, potentially guiding targeted interventions to enhance cattle reproductive health and productivity. Further exploration into the complex interactions between rumen microbes and host physiology could lead to more refined strategies in cattle management and breeding, optimizing both animal welfare and agricultural efficiency.

## 5 Ethics Statement

All experimental procedures were approved by the Committee of the Care and Use of Experimental Animals at the Graduate School of Bioagricultural Sciences, Nagoya University.

## 7 Author Contributions

S.O., J.M., Y.M., S.N., S.N., and V.P.R. conceived, designed, and supervised the project. YU and VPR conducted the field sampling and molecular experiments. VPR conducted the informatics work. VPR wrote the initial draft of the manuscript. All authors read, edited, and approved the final manuscript.

## Acknowledgments

The authors would like to acknowledge Ms. Kinuyo Yamazaki, Mr. Fumitaka Yoshimura, and Mr. Yoshiki Kono for the technical assistance and animal care. Mr. Michimasa Oguri for cooperation in the collection of rumen fluid samples.

## 9 Supplementary Material

## Data Availability Statement

The datasets generated and/or analyzed in this study are available at the European Nucleotide Archive database under the accession number PRJEB76958.

